# Not All Experimental Questions Are Created Equal: Accelerating Biological Data to Knowledge Transformation (BD2K) via Science Informatics, Active Learning and Artificial Intelligence

**DOI:** 10.1101/155150

**Authors:** Simon Kasif, Stan Letovsky, Richard J. Roberts, Martin Steffen

## Abstract

Pablo Picasso, when first told about computers, famously quipped “Computers are useless. They can only give you answers.” Indeed, the majority of effort in the first half-century of computational research has focused on methods for producing answers. Incredible progress has been achieved in computational modeling, simulation and optimization, across domains as diverse as astrophysics, climate studies, biomedicine, architecture, and chess. However, the use of computers to pose new questions, or prioritize existing ones, has thus far been quite limited.

Picasso’s comment highlights the point that good questions can sometimes be more elusive than good answers. The history of science offers numerous examples of the impact of good questions. Paul Erdős, the wandering monk of mathematical graph theory, offered small prizes for anyone who could prove conjectures he identified as important (*1*). The prizes varied in cash amounts based on the perceived complexity of the problem posed by Erdős.

Posing technical questions and allocating resources to answer them has taken on a new guise in the Internet age. The X-Prize foundation (http://www.xprize.org/) offers multi-million dollar bounties for grand technological goals, including goals for sequencing genomes or space exploration. Several companies provide portals where customers can place cash bounties on educational, scientific or technological challenges, while potential problem solvers can compete to produce the best solutions for these problems. Amazon’s Turk site (https://www.mturk.com/mturk/welcome) links people requesting performance of intellectual tasks to people willing to work on them for a fee. Such crowd-sourcing systems create markets of questions and answers, and can help allocate resources and capabilities efficiently.

This paradigm suggests a number of interesting questions for scientific research. In a resource limited environment, can funds and research capacity be allocated more efficiently? Can knowledge demand provide an alternative or complementary mechanism to traditional investigator-initiated research grants?

The fathers of Artificial Intelligence (AI) and Herbert Simon in particular envisioned the application of AI to Scientific Discovery in different forms and styles (focusing on physics). We follow on these early dreams and describe a novel approach aimed at remodeling of the biomedical research infrastructure and catalyze gene function determination. We aim to start a bold discussion of new ideas aimed towards increasing the efficiency of the allocation of research capacities, reproducibility, provenance tracking, removing redundancy and catalyzing knowledge gain with each experiment. In particular, we describe a tractable computational framework and infrastructure that can help researchers assess the potential information gain of millions of experiments before conducting them. The utility of experiments in this case is modeled as the predictive knowledge (formalized as information) to be gained as a result of performing the experiment. The experimentalist would then be empowered to select experiments that maximized information gain if they wished, recognizing that there are frequently other considerations, such as a specific technological or medical utility, that might over-ride the priority of maximizing information gain. The conceptual approach we develop is general, and here we apply it to the study of gene function.

## What are the best genes to study? Prioritizing gene function research

The newly established initiative in Biological Data to Knowledge Transfer (BD2K) (http://commonfund.nih.gov/bd2k/index) aims to develop transformative approaches to maximize the integration and utility of Big Data into biomedical research. While a conservative view might suggest that knowledge is just a set of experimentally validated facts, modern sciences are often driven by predictive models that organize facts, which cumulatively enhances our ability to predict behavior of complex physical or biological systems.

The field of genomics provides an interesting case study for the allocation of research resources to BD2K. High-throughput sequencing has led to the discovery of a plethora of new genes, many of which are poorly understood. Each gene gives rise to a set of questions: what is its function (*2*, *3*)? If it codes for an enzyme, what substrates does it act on? If it codes for a regulator, what does it regulate (*4*)? What is its role in the life of the organism (*5*)? Is the function maintained across conditions and across species? How did it evolve? Does the gene have an important clinical relevance such as contributing to antibiotic resistance (*6*) or cancer (*7*)?

A fairly typical protein cluster contains hundreds of evolutionary related proteins. While each one of the questions above can produce a unique answer for each gene in this cluster, we will never be able to experimentally test all possible proteins. If there is a hypothesis or a prediction about the biochemical function performed by any of the proteins in this cluster, which specific experiments should we conduct, among hundreds of possible options? While numerous bioinformatics systems offer an ability to predict the function of a protein, none answer the question: “What is the best experiment to perform?” Should the experiment be exclusively driven by curiosity, interest in a particular bacterial organism or specific properties of the protein family? Is there a general paradigm that can be deployed in guiding the choice for experiments?

We describe one solution to this question in the context of bacterial gene function determination. The impact of the biochemical characterization of newly sequenced bacterial enzymes can be very high. Two recent “Breakthroughs of the Year” have been Optogenetics (Nature, 2010) and CRISPR (Science, 2015), both of which rely on bacterial enzymes! As is well known, optogenetics relies on light-regulated ion channels, and genome editing with CRISPR relies on sequence-guided nucleases. Currently on NCBI (*8*) and other protein databases, there are protein clusters that contain mixtures of nucleases and proteases. However, we are not able to discriminate which is which, in part because the essential experiments have not yet been conducted. There are hundreds of similar examples - an immense knowledge mine of possible technology drivers in the form of bacterial proteins used to produce energy, catalyze natural product synthesis, perform sensory activities, and which could play a key role in synthetic biology applications.

## Does the power-law “curse” inhibit knowledge growth in molecular biology?

Although many high-throughput technologies for generating data related to aspects of gene function have been developed (expression levels, interacting molecules, phenotype, *etc*.), most experiments specifically on the molecular function of a hypothetical protein are answered using low throughput methods. How does the scientific community collectively decide which genes and gene families to focus on for function elucidation? There are thousands of proteins clusters that lack any or specific annotation. Each of these clusters may contain hundreds of proteins. Which proteins are the “best” to test?

Publication statistics show that the allocation of research attention across genes is far from uniform, and instead follows a power law distribution (*9*, *10*) In other words, most genes have zero or few citations, and a very few genes are associated with very large number of citations (previously observed for human genes). Power law distributions often arise as a consequence of preferential attachment dynamics (*10*) in which “the rich get richer.” In the scientific literature, genes discussed in a lot of publications tend to generate more manuscripts discussing them. Many factors might contribute to this apparent bias: the more heavily published genes may be involved in more medically important diseases; they may have been discovered earlier, and so have had more time to accumulate publications; they may be “hubs” in cellular networks that inherently affect more biological processes, or they may be more amenable to study using the currently available methods. By far the most popular gene product, measured by number of publications, is the tumor suppressor p53, for which many of the above criteria might apply. In bacteria, the ten most popular genes are recA, rpoA-rpoD, rpoS, dnaK, ftsZ crp and rne, which are also highly conserved e.g. the rpo genes encode RNA polymerases.

However, the bias in favor of particular genes may be self-reinforcing. Peer review of grants and publications favors the currently fashionable and familiar genes, thereby encouraging additional allocation of resources to them. The result can be a self-perpetuating “power law curse”: a fashion-driven science, with a biased allocation of attention and resources that deflects attention away from the least characterized genes. The distribution of publications per *E. coli* gene at left is roughly linear on a log-log plot, characteristic of a “power-law curse”. Such distributions are often due to preferential attachment, also known as the “rich get richer” or “Matthew” effect (*11*). The right plot shows a positive power law relationship between publications per gene and protein cluster size, showing that more conserved and/or widespread proteins tend to have more publications.

## COMBREX: A Computational Bridge to Experiments

The COMBREX project (http://www.combrex.org/, http://combrex.bu.edu), sponsored by the National Institutes of Health, is a transformative effort designed to accelerate the acquisition of gene function knowledge in bacteria by a novel funding mechanism and Artificial Intelligence. COMBREX awarded modest funding for experimental tests of gene function, that were prioritized in parallel by (*1*) explicit, theoretically grounded criteria, or (*2*) by a selection of practical, biomedically motivated criteria (*e.g.* essentiality, conferring antibiotic sensitivity, *etc.*) (*12*, *13*). COMBREX, an acronym for “COMputational BRidges to EXperiments,” forged links among computational and experimental communities. The project was built on many years of experience gained by both groups, doing transformative work on biochemistry, genome sequencing, gene discovery, and gene function prediction, coupled with an organizing call for community action (*14*). The project initially focused on biochemical, molecular function (*e.g.* enzymatic activity), but similar approaches can be developed for other biological function annotation classes (*e.g.*, pathway membership or phenotype). Similar to other resources, such as Gene Ontology (GO) or model organism databases, COMBREX has amassed a database of bacterial gene function assignments from both experimental evidence and computational predictions, along with information on the quality and type of evidence supporting each assignment (*15*). Unique to COMBREX-DB is the influence and implementation of revolutionary AI proposals made more than 40 years ago to develop intelligent systems able to guide the Scientific Discovery process (*16*). The database and the webserver are intended to function as a Science Hub to coordinate activities required for the generation and tracking of microbial gene function knowledge. The overall effort seeks to drive this database towards correctness and completeness by developing the four following principles.

### Computable Function Descriptions

64% of bacterial gene functions are described only by narrative text gene descriptions, which among other issues, rarely describe the specific substrates of a protein. It can be challenging for a computer to determine when two textual function descriptions that are not worded precisely identically are describing the same function, or when one description adds additional details to an otherwise identical description. Formalization of gene descriptions using controlled vocabularies such as the Gene Ontology Molecular Function hierarchy (*17*), the Enzyme Commission numbering scheme, and explicit reaction descriptions is essential for enabling automated reasoning over functions, including logical (e.g. equality or similarity), statistical (e.g., accuracy assessment) and biological (e.g. metabolic reconstruction) comparisons and analysis of predictions. COMBREX is developing and encouraging the community to produce strategies to increase the extent of structured annotation using both automated and manual curation methods. Such systems (translating text to structured descriptors) are under development in multiple bioinformatics and natural language processing groups and will continue to improve the formal knowledge of gene function (*13*).

### Traceable Evidence

The evidentiary basis of many gene function assignments is either poorly recorded or missing. Although some gene function assignments are based directly on experiments, the vast majority are inferred from sequence similarity: the function assigned to a protein is based on an experiment carried out on a protein with some degree of homology, typically from another species. The expected accuracy of such propagated or predicted biochemical function predictions often depends on the degree of sequence similarity between the homologous proteins, but details are important. Single amino acid changes in catalytic residues can alter an enzyme’s function, while other times, numerous substitutions in a protein may have no effect. There is mounting evidence (*18*) that propagation of function by sequence similarity is error prone and creates a significant mis-annotation problem that is exacerbated by the number of genes added daily to NCBI and other genomic databases.

Surprisingly, COMBREX determined that less than 1% of bacterial function assignments to genes are supported by experimental evidence that can be ***explicitly traced*** to a publication (see Figure 2). Often it is unclear which function assignments are based on experimental evidence rather than homology, which protein sequence was used in an experiment, what the publication is, or how assignments were made. This lack of traceability, or provenance, makes it difficult to detect and correct errors; it can even lead to self-supporting cycles of evidence, as functions are propagated back to their starting points, often across databases. To address this, COMBREX has developed a color code for annotating evidence quality: gold for experimentally validated functions that have been manually curated, green for functions that have been experimentally validated but await curation, blue for computationally predicted functions, and black for genes with no functional information. The scheme is borrowed from ranking the degree of difficulty in downhill skiing, and is aimed to suggest the challenge level in experimental testing. In addition to the COMBREX database (*15*), this color scheme has recently influenced the SMART BLAST engine built by NCBI (*8*). Evidence propagation trails are either recorded or **predicted** in the COMBREX database to provide traceability. COMBREX color codes are an experiment centric refinement of the GO evidence codes, but COMBREX is the first to implement a simple (straw man) algorithm to predict an experimental source for a predicted annotation to drive an integrated process of evidence tracking and improvement. This is done simply by BLASTing to all experimentally determined proteins and providing a link to the best matches.

### Computational Predictions

In its first iteration, the COMBREX project assembled a computational consortium to develop improved gene function prediction methods. It included researchers from structural biology, machine learning, metabolic modeling, systems biology, and comparative genomics. Results from such a wide range of predictive methodologies have not previously been integrated in a common repository. The groups used a variety of algorithmic approaches to predict gene functions, including pairwise sequence similarity (*19*), profile-based approaches (*20*), module-based approaches (*21*), evolutionary modeling (*22*, *23*), functional linkage (*2*, *24*), genomic context (*25*, *26*) gap filling in metabolic networks (*27*), genomic role filling (*28*), structural remote homology modeling, threading (*29*), protein-ligand docking (*30*, *31*) and combinations of these (*32*). In addition to publishing predictions generated by consortium members, COMBREX solicited predictions from the larger computational community, with the goal of serving as a central archive of predictions. Predictions can be readily uploaded into COMBREX-DB. Function prediction is a relatively well established area in computational biology with many competing algorithms and benchmarks (*33*). There are many benefits to such competitions, which have generally shown that no single algorithm is likely to be best in biochemical function prediction for every type of enzyme. In most cases, specialized protein sequence-structure-function approaches that take into account specialized evolutionary and functional constraints are needed to produce the best predictions of putative substrates. COMBREX will continue to collaborate broadly with multiple specialized teams and will aim to fund small computational projects that specialize general computational approaches (*e.g.* SIFTER (*34*)) to a specific protein families.

### Prioritized Targets for Experimental Validation

At present writing ~2.6 million bacterial genes in the COMBREX database have computational predictions, while fewer than 1% have documented experimental validations. In the absence of reliable high throughput methods for gene function testing, there are many more predictions than can be experimentally tested on any reasonable timescale using current methods. This suggests that for the foreseeable future, knowledge of gene function is going to be fundamentally probabilistic, that is, inferred computationally from known experiments and analysis using methods of varying reliability. Can we assess probabilistic knowledge by some formal measure and use it to guide the selection of experiments? Can we prioritize gene targets in a way that integrates the desire to focus on practically important genes with the goal of maximizing the overall growth in knowledge?

COMBREX implemented a proof-of-concept, automated recommendation system to help experimentalists identify proteins to include in a planned experiment on a given protein family. Recommendations take into account a number of factors, but a key consideration is information gain; i.e., the expected impact of a validation on the annotation of homologous or functionally linked (*24*, *35*) proteins. Information gain can take different forms depending on the method or model used for gene function prediction. The simplest measure is the number of proteins in a protein family. This metric will bias gene exploration towards more conserved and widespread genes, genes that are essential for living organisms or specific species. Alternatively, for function prediction methods that report probabilities with their predictions (*24*), the information gain from an experiment can be quantified as the reduction in the estimated probability of prediction error, summed across all predictions. Similarity-based prediction methods can estimate prediction probabilities based on the BLAST distances to experimentally validated proteins, if one assumes that functional divergence is related to sequence divergence. When functional selection is present, positive selection drives a higher rate of sequence divergence away from a cluster center.

In the initial COMBREX-DB, we implemented a novel yet simple feature that attempted to guide experimentalists towards validations of “consensus” proteins. The logic supporting the feature can best be illustrated by considering the example of a large cluster of proteins that has several hundreds of proteins but no experimental annotation. A simple histogram provides an estimate of how many proteins need to be tested to sample the evolutionary span and functional divergence found in a protein family. This preliminary work was implemented for approximately 500,000 protein clusters annotated by NCBI in COMBREX-DB (*12*, *13*, *15*, *36*-*38*). The proteins are ranked based on the average phylogenetic tree distance to all other proteins in the cluster, and a ranked list of all proteins in the family are prioritized by shortest average distance. For some protein families characterization of a single protein might be a good overall representation of the protein family’s activity (Fig 1, Panel A) which shows no underlying structure to the cluster. If one were to test a second protein from that family, one might pick an outlier, a protein with a large average distance from the centroid. If the activity is found to be essentially similar, it increases the confidence that the family is likely to have relatively homogeneous function. We built this into the original version of COMBREX displaying the histograms based on multiple alignments and evolutionary trees, and returning a ranked list of members within a protein family (Fig 3, bottom).

**Figure 1:**
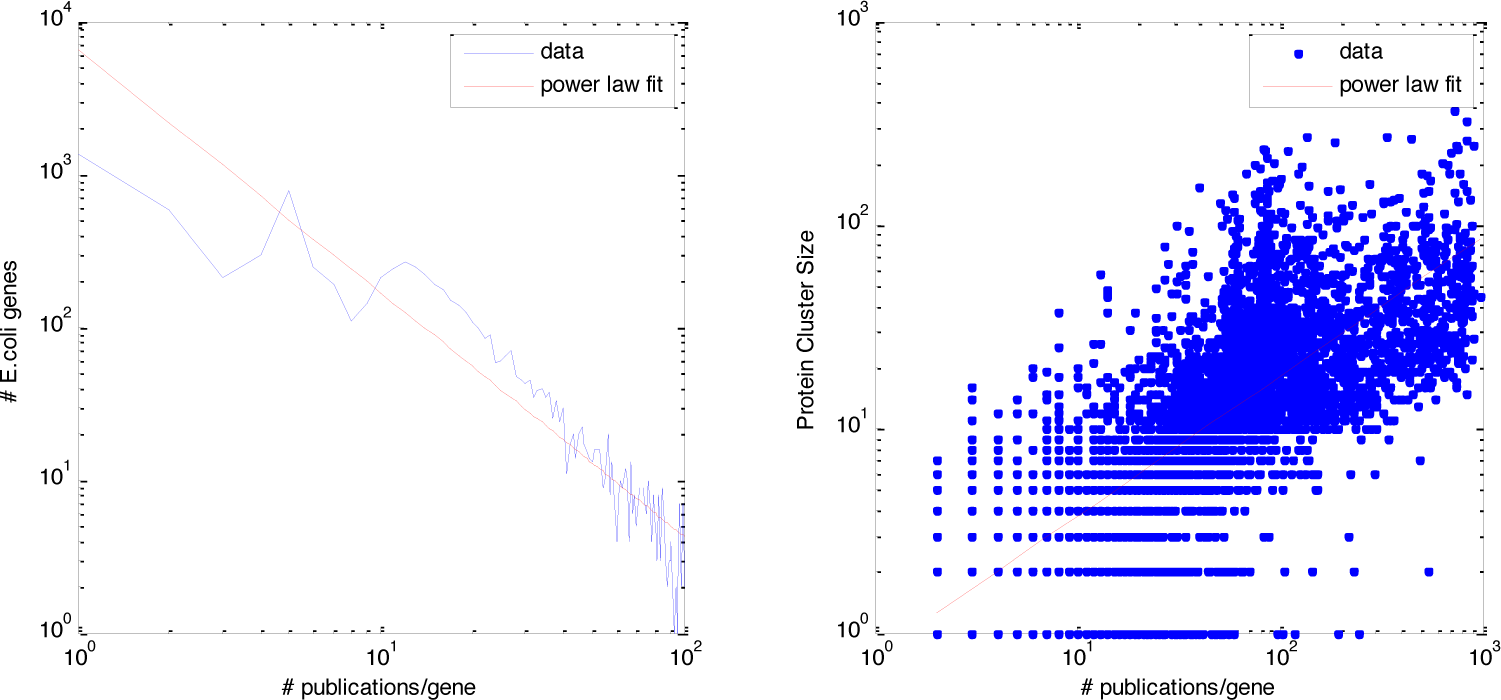
The distribution of publications per *E. coli* gene

**Figure 2:**
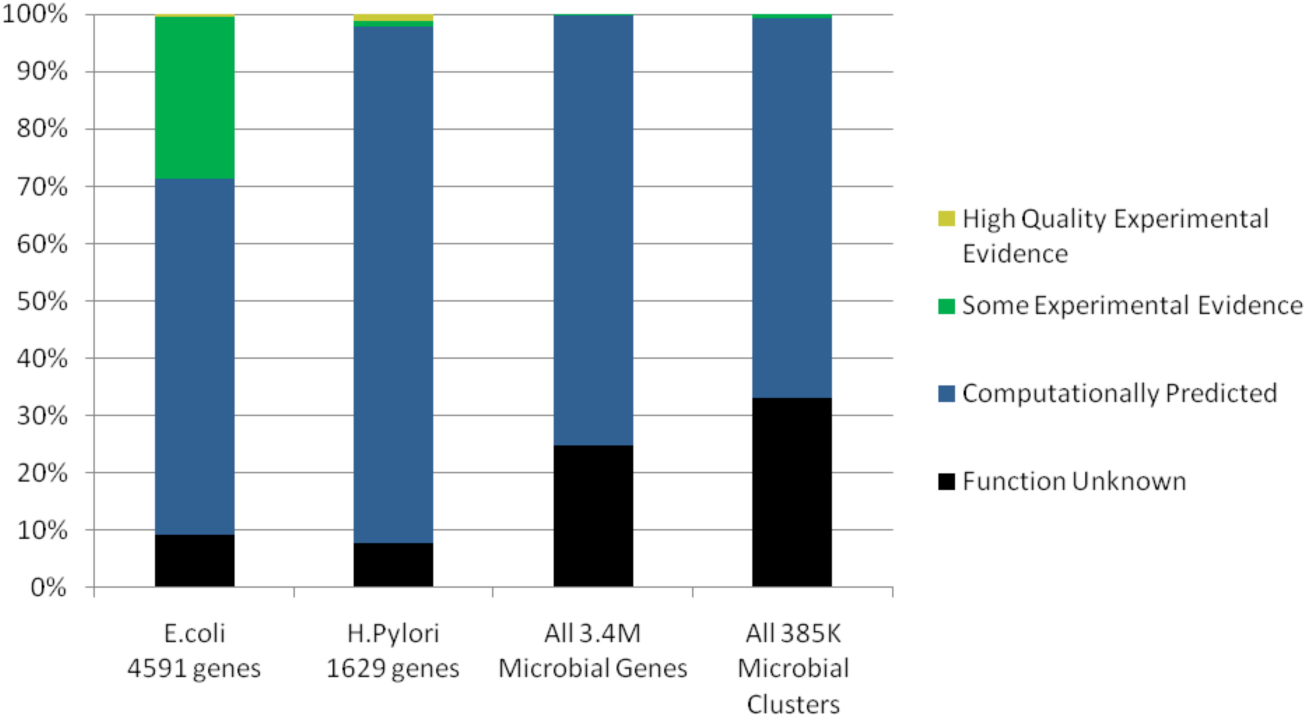
Function annotation rates for bacterial genes, using the COMBREX evidence color codes. Fewer than 1% of bacterial sequences have high quality experimental evidence of function.

**Figure 3:**
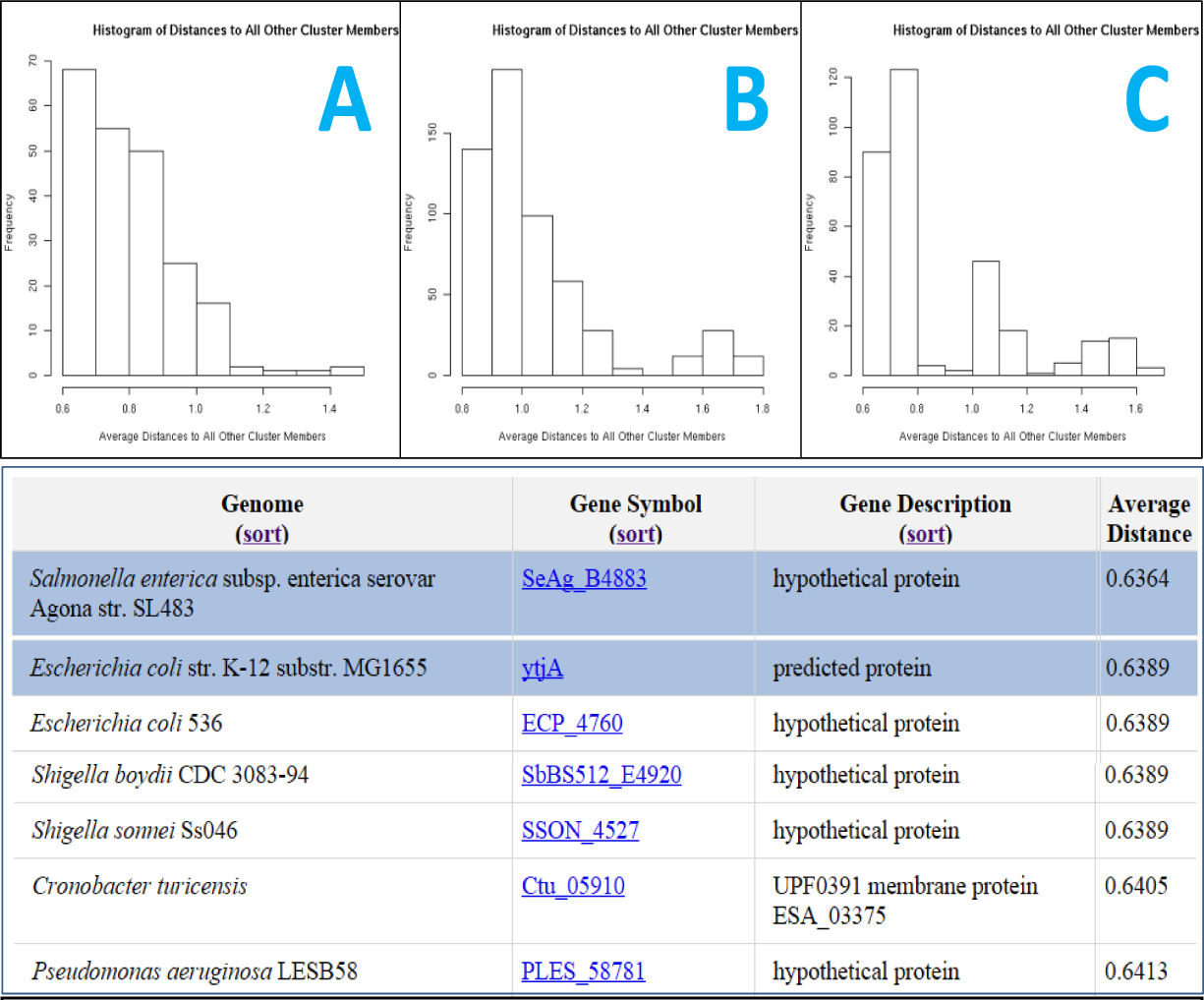
Histograms of tree distances to guide experimental selection.

When the histograms become multi-modal (Fig 3, Panel B - 2 subclusters; Fig 3, Panel C - 3 subclusters) we can be reasonably certain that characterization of a single protein from these would not be adequate, and that testing multiple proteins would be required. This type of information is not available on any existing database and we feel could have a significant impact if available to biologists.

There are many other possible considerations for the prioritization of experiments testing protein function. An experimentalist might be interested in a specific organism, pathogenic bacteria, proteins that confer drug resistance or susceptibility, proteins that have solved structures, proteins with homology to human proteins, or proteins with PFAM domains of unknown function. Experimentalists were able to consider all of these selection criteria, which were incorporated into the first prototype of COMBREX working closely with many collaborators (*13*).

### Next Generation Informatics: Using Active Learning and Information Gain from Experiments

The computational model used to generate a prediction can affect the rankings of experimental prioritization, *e.g.,* if a residue is predicted to be catalytically important, then experiments should target proteins that contain variations in that residue. For widely used probabilistic predictive systems, including profiles produced from multiple alignments, well-established frameworks can be used as starting points. Before an experiment is conducted each protein is associated with a random binary variable X that quantifies the probability whether the protein has particular function or not (*e.g*. nuclease or protease). Without any experimental evidence or models we can only use the relative empirical frequencies to estimate these probabilities. As one very simple example we illustrate the fairly simple mathematics needed measure the probabilistic knowledge learned from a single experiment as it pertains to all related inferences we can make based on the new data.

The entropy of each binary protein class variable is defined as usual: H(x) = - Sum_i_ p(x_i_) log_2_ p(x_i_)

The reduction in the entropy, E, of the prediction is the Information Gain (IG) obtained by learning the class label of the protein. The IG is *(E before experiment – E after experiment)*. The experimental testing of a single protein will cause a cluster-wide decrease in entropy that is the sum of the decreases in entropies of all proteins that are functionally linked (by any method) to the tested protein. The value of the overall decrease in entropy is the information gain (IG) associated with any planned or conducted experiment.

The KL-divergence is a measure of how different two distributions P and Q are:

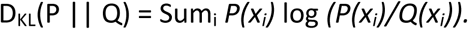

While KL-divergence is formally not a distance, it can be used as a proxy to quantify the reduction in entropy of all predicted variables (functional labels) in the protein cluster given a single experiment or more. More formally, one would measure the sum of the divergences between the posterior distributions (after an experiment) and the prior distributions for all proteins in a cluster as the IG of an experiment on every protein and prioritize experiments based on these scores. This formally quantifies what we learned or might learn from an experiment.

To make our proposal concrete we describe a specific implementation of this principle using protein profiles (*39*). A protein profile is really a simple multinomial model that specifies the probability of any residue occurring at a given position in the profile (*40*-*42*). Given a profile, we can determine the log-likelihood that a protein belongs the functional family (*i.e.* matches the profile) by summing the log probabilities at each position and normalizing by the logs of prior probabilities. Profiles have been generalized to Hidden Markov Models (HMM), and disseminated using PFAM (*39*). HMM-style models are routinely used for inferring functional labels (*e.g*. TIGRFAM). We will describe our information gain principle for a position-specific scoring matrix, PSSM, here, but a similar approach is readily extended to HMM models or their various generalizations in the form of graphical models proposed by us and others.

Assume we have computed a PSSM profile for a family of proteins. This means that for each protein in the protein cluster or family we can compute a probabilistic score of a protein being a member of this family (*39*, *43*). This probability is computed for each column in the PSSM matrix and then the log-probabilities of each column are summed across the entire alignment (*39*). Let us refer to this probability as *P(Prot | PSSM)* (*39*, *43*). We compute the entropy of this prediction for both single proteins and the entire cluster (by summing entropies). Under simplifying but realistic assumptions, one can show that the best single experiment is the “center” of the cluster defined formally by the protein X* such as: X* = arg max P(X | PSSM).

One can generalize this approach to non-homology functional linkage and prediction. Probabilistic Functional Linkage Network Graphs (PFLG), introduced by us and others (*44*-*46*) to formalize and systematically implement Guilt-By-Association inference in Functional Linkage Networks (FLNs) (previously introduced by David Eisenberg, Ed Marcotte and Matteo Pellegrini). FLNs (*26*, *32*, *35*, *47*-*49*) employ nodes to represent proteins and edges to represent evidence of common function. Edges can be weighted to represent the strength of that evidence (*3*, *50*). Edges may come from protein-protein interaction (PPI) data, correlated gene or protein expression profiles, correlated term usage in literature, or others (*51*, *52*). The most popular resources for FLNs are available in STRING (*53*) and VISANT (*54*) or by using methodologies reported in (*52*, *55*-*57*). Genomic context methods provide information complementary to sequence homology such as phylogenetic profiles (*58*, *59*), chromosomal gene clustering (*60*-*62*), and protein fusion (*63*-*65*), metabolic networks (*20*) and others (*26*, *58*-*63*, *65*-*67*).

There are several natural ways to define maximally informative proteins in FLNs. Kleinberg and coworkers (*68*) introduced the theoretical foundation for this network problem for issues associated with the internet, called influence networks. Intuitively, all measures related to high degree or centrality in FLNs are good straw-man heuristics for choosing graph “centroids” as best experiments (*69*). Choosing K-best experiments becomes computationally intractable (similar to warehouse location problems in distribution networks), but simple heuristics for graph partitioning usually work.

Other things equal, prioritizations might also be biased towards biomedically important or specific bacterial model organisms. Other criteria include medically-important gene phenotypes such as essentiality, pathogenicity, antibiotic resistance, biofilm formation, and growth. These considerations may not break the existing power-law “curse” or replace it by another (*9*), but we emphasize that these computed prioritizations are intended as suggestions to the experimental scientist, and are not mandated by the system or required in order to participate in COMBREX. There will always be a place for creativity and curiosity in the choice, design and execution of experiments; COMBREX’s prioritizations simply help raise awareness of information gain and medical relevance for experimentalists to take into account in the search for gaps in gene function knowledge.

These experimental prioritization criteria complete a cycle of increasing knowledge (Fig. 4). In the field of machine learning, a cyclic process that attempts to optimize the choice of the next experiment to maximize information gain or predictive accuracy is termed *active learning (*70*)*. COMBREX appears to be the first implementation of a science informatics systems using a community-based active learning paradigm, coupling the vast capability and expertise of human research communities with integration of predictions and web-based communication, thereby creating an active learning loop for bacterial genomics. However, while it is too early to say if this approach will result in a more productive utilization of research capital than more traditional funding mechanisms, COMBREX provides a rejoinder to Picasso’s critique of the limits of computers in providing answers but not questions. It is hoped that these approaches, as implemented in COMBREX, will catalyze an increased rate of growth in biological knowledge, while enhancing the bridge between experimentalists and bioinformaticians.

**Figure 4:**
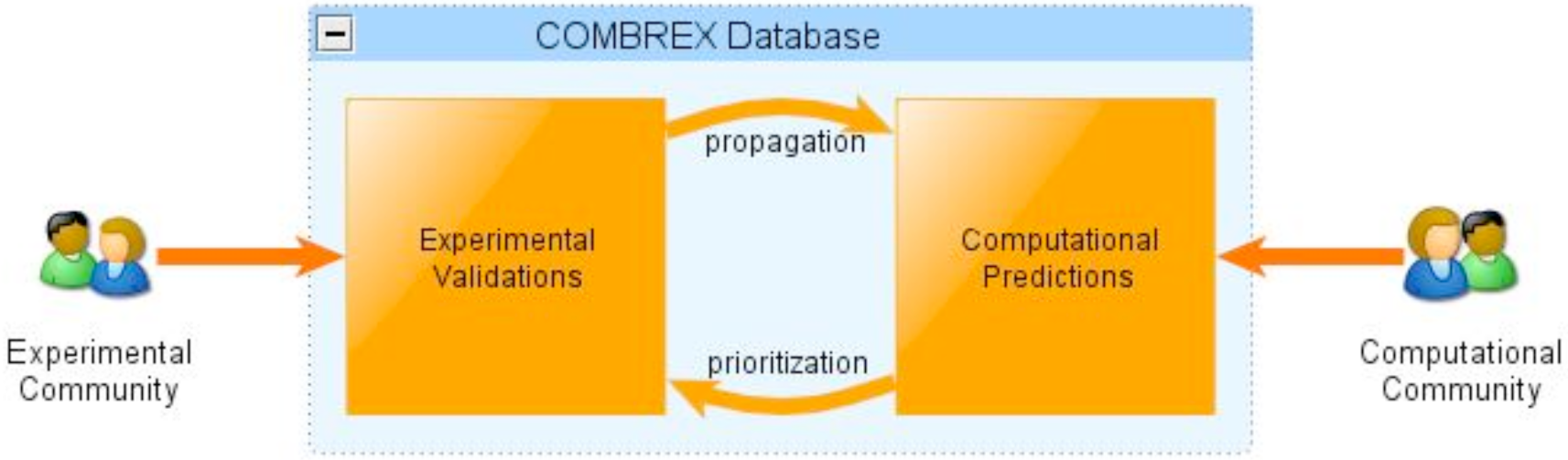
The COMBREX Active Learning Loop. The database acts as a bridge between the computational and experimental communities, storing computed function predictions and experimental validations. Propagation of experimental results through predictive inferences yields new predictions, while intelligent prioritization of predictions leads to new “most informative” experiments.

Last but not least we want to highlight three philosophically important ideas intimately intertwined with our proposal. Computational Biology is a fast evolving field that made enormous contributions to life sciences. The advent of genome sequencing and other high-throughput omics technologies shifted bioinformatic emphasis towards the development of specific technical algorithms to interpret the rapidly growing BIG data. The current focus on comparing and improving the interpretation algorithms are not sufficient on their own to produce biological knowledge. It would be beneficial to shift some emphasis to the development of systems that manage, monitor and track the entire Science Informatics and Discovery process, including driving the “most informative” experiments that will catalyze growth in knowledge, track provenance, remove redundant research and improve reproducibility. In order to most effectively support the BD2K pipeline, systems need to also track knowledge gaps, not just experimental data and predictions. The revolutionary “Robot Scientist “proposal (*71*) proposed an “extreme” solution, leading to robot scientists performing the “best” experiments replacing experimentalists. The COMBREX model is a compromise supporting an Amazon Turk – Citiizens Science model, which builds a system that tracks knowledge, knowledge gaps and provides AI based active learning strategies to help guide informative experiments. If computers can recommend movie selections and stock picks, they can also recommend experiments.

A second point is to “liberate” computational biology, and position bioinformatics as a driver of science helping design experiments. This is in contrast to the existing pipeline in which experimentalists generate data, whereas bioinformaticians and biostatisticians devise algorithms to interpret it (*24*, *33*, *53*, *72*, *73*). In the approach highlighted by COMBREX, computational systems generate prioritized experimental questions, and biologists are funded to experimentally test specific regions in the “gaps” of biological knowledge.

The COMBREX webserver and database created a broad computational science informatics infrastructure, envisioned by early AI, and specialized to gene function prediction and management of the scientific discovery process, by including predictions, prioritizations, management of bids, and other activities. COMBREX advances the concept of “Citizen Science.” In addition to reaching out to experimental laboratories, COMBREX also worked with undergraduate classroom laboratories with expert supervision (*74*).

As a proof of concept COMBREX also established a novel micro-granting mechanism to support experimental validations of prioritized gene functions, a major focus for COMBREX. Experimentalists can search the database for prioritized gene function predictions in their areas of expertise, and submit brief proposals to experimentally validate those predictions. During the initial proof of concept period, proposals went through a rapid peer review, grants from a few thousand to a few tens of thousands of dollars were awarded to experimentalists to test predictions for a small number of genes including prioritized targets. Awards included grants to foster educational activities for experimental testing by undergraduates under expert supervision in classroom or research labs. As of this writing, over 150 genes have had functions tested in response to many bids from experimentalists.

The COMBREX proposal was presented at a Common Fund Workshop organized by the office of the NIH Director to solicit particularly transformative and broadly useful ideas in biomedical sciences (across all funded disciplines at NIH). It was presented as a package that included other Citizen’s Science proposals, such as the Connectome project (*75*) and the Foldit Protein Folding game (*76*). So far, these proposals have not generated a robust Citizens Science infrastructure to advance biomedical sciences. Recently, several prominent scientists have called for related ideas and paradigm changing transformations. We hope that the renewed interest in AI, coupled with the experience gathered in Bioinformatics, will create a movement in the community towards recognizing this unique new opportunity in biological sciences, and lead to a new paradigm in which AI and computational biology can help drive biology, in part by enabling a faster and more robust growth in critical biomedical knowledge.

## Acknowledgements

We thank all members of the COMBREX consortium (www.combrex.bu.edu) that include participants, advisory board members, and numerous scientists who supported the effort, the COMBREX workshops and proposals. We thank Yi-Chien for Figures 1 and 2. We thank the NIH stimulus program for funding parts of this effort.

